# Catalyzed Synthesis of Zinc Clays by Prebiotic Central Metabolites

**DOI:** 10.1101/075176

**Authors:** Ruixin Zhou, Kaustuv Basu, Hyman Hartman, Christopher J. Matocha, S. Kelly Sears, Hojatollah Vali, Marcelo I. Guzman

## Abstract

How primordial metabolic networks such as the reverse tricarboxylic acid (rTCA) cycle and clay mineral catalysts coevolved remains a mystery in the puzzle to understand the origin of life. While prebiotic reactions from the rTCA cycle were accomplished via photochemistry on semiconductor minerals, the synthesis of clays was demonstrated at low temperature and ambient pressure catalyzed by oxalate. Herein, the crystallization of clay minerals is catalyzed by succinate, an example of a photoproduced intermediate from central metabolism. The experiments connect the synthesis of sauconite, a model for clay minerals, to prebiotic photochemistry. We report the temperature, pH, and concentration dependence on succinate for the synthesis of sauconite identifying new mechanisms of clay formation in surface environments of rocky planets. The work demonstrates that seeding induces nucleation at low temperatures accelerating the crystallization process. Cryogenic and conventional transmission electron microscopies, X-ray diffraction, diffuse reflectance Fourier transformed infrared spectroscopy, and measurements of total surface area are used to build a three-dimensional representation of the clay. These results suggest the coevolution of clay minerals and early metabolites in our planet could have been facilitated by sunlight photochemistry, which played a significant role in the complex interplay between rocks and life over geological time.

## Introduction

One of the major scientific questions that remains uncertain is how the origin of life occurred.^1^ Within the prebiotic chemistry context of this interdisciplinary problem, recent work has shown the potential of ZnS promoted photocatalysis to harvest sunlight energy into chemical bonds.^2^ The mechanism of semiconductor promoted photochemistry may have played a major role in cycling small organic compounds essential for the origin of life.^3^ For example, illuminated ZnS has successfully driven several reactions of the reverse tricarboxylic acid cycle (rTCA),^3^ which is central to metabolism. Our latest work explored in detail the photoreduction of fumarate to succinate on the surface of ZnS,^4^ which proceeds with a 95% yield.^5^ The synergistic interaction between sunlight, photocatalysis, and organic acids in the prebiotic Earth could have promoted reactions otherwise unviable, providing the foundation for present metabolism.^3^

Within the origin of life framework, a separate but relevant problem is the potential role of clay minerals, strong adsorbents of polar organic molecules,^6^ to facilitate abiogenesis.^7^ The catalytic power of clays can promote the polymerization of biomolecules and the conversion of fatty acid micelles into vesicles.^7^ Indeed, the process of clay formation is key for developing an understanding of the possible roles of these minerals in the origin of life.^8^ Recent work has proven the crystallization of saponite clays can proceed easily in only 20 h under relatively mild conditions in the presence of urea^9^ as a catalyst. Interestingly, nowadays abundant oxalic acid, an endpoint oxidation product for all organic matter exposed to environmental oxidizers before their final conversion into formic acid and CO_2_,^10–12^ successfully substituted urea in the synthesis of saponite clays.^13^ The role of oxalate as a chelating agent for aluminum atoms in the octahedral state has been previously studied.^14^ Bare Al^3+^ has been proposed to retain the hexacoordination for crystalizing the phyllite structure when its complex with oxalate is decomposed in the presence of silicate.^14^

Despite the previous knowledge in both fields, the unanswered question remaining is whether a relevant clay formation mechanism could have proceeded catalyzed by photogenerated central metabolites^3^ of the rTCA cycle. In this work, succinic acid (and other organic acids) is used as a probe from the intermediates of the rTCA cycle to catalyze the synthesis of sauconite, a model zinc clay mineral from the smectite family. The work hypothesizes that succinate accommodates in the interlayer space after catalyzing the incorporation of Al^3+^ into the precursor soluble gel, dominated by SiO_2_, to create the tetrahedral layer. The work shows that sauconite crystallization is related to the ability of succinate to chelate bare Al^3+^. Syntheses of sauconite are performed under variable initial concentration of sodium salts of carboxylic acids (e.g., [succinate]_0_), pH, and temperature, to monitor during 20 h the nucleation process. Succinate is shown to catalyze the reaction of Al^3+^ with silicic acid, zinc and sodium compounds, yielding an authentic 2:1 clay mineral of the smectite group. The synthesis of Zn-clay material proceeds at low temperature (<100 °C) and ambient pressure (~1 bar) as demonstrated from the structural characterization displaying the early stages of the synthesis. This characterization is performed by a combination of techniques including: 1) Powder X-ray diffraction (XRD), 2) diffuse reflectance infrared Fourier transform (DRIFT) spectroscopy, 3) conventional transmission electron microscopy (TEM), and 4) cryogenic TEM (cryo-TEM).

## Results and Discussion

The bottom-up synthesis of sauconite, with a theoretical formula Na_1.2_Zn_6_{Si_6.8_Al_1.2_}(O_20_)(OH)_4_•*n*H_2_O,^6,15^ is performed from a silicic acid gel in contact with Zn^2+^, Al^3+^, and succinate^2-^ ions. The synthesis can be easily altered to vary the components integrated into the structure. Aiming at enhancing nucleation and to accelerate the formation of clay minerals, we designed a set of experiments to optimize the synthesis of a model sauconite. To study the role of succinate, a central metabolite, as a catalyst promoting clay formation, we made several modifications to the gel. In the first set of experiments we varied the concentration of sodium succinate (Figure 1). Alternatively, the nature of the organic salt is investigated by substituting succinate with sodium salts of formic acid, acetic acid, oxalic acid, and malic acid (Supplemental Fig. S1-S4 online). In the second set of experiments, we performed the synthesis under variable temperature. In the third set of experiments, the initial pH was varied in the range 6-14.

**Figure 1.**
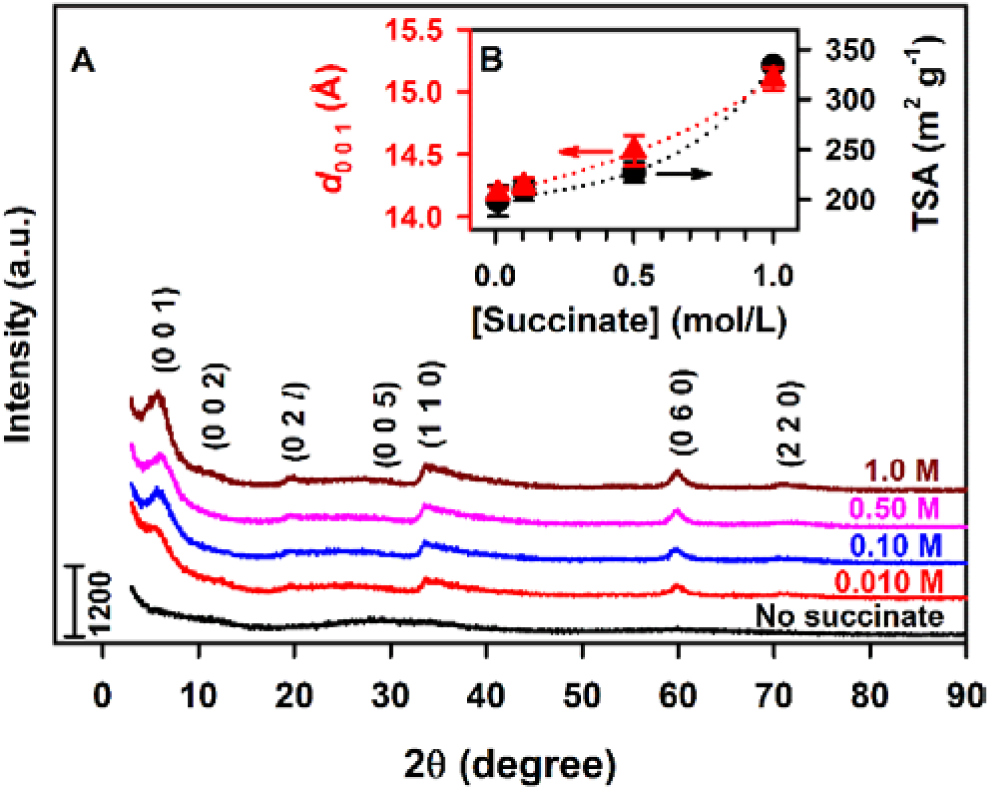
Powder XRD and TSA characterization of sauconite synthesized at pH_0_ 6.3 and 90 °C during 20 h under variable succinate concentration. (A) (A) XRD diffractograms for [succinate] listed above each trace. Numbers, e.g., (0 0 1), indicate basal reflections of the identified phases. (B) The layer to layer distance for 2:1 sauconite (*d*_0 0 1_, red solid triangle) and TSA (black solid circle) for traces in (A) vs [succinate].

### Effect of Varying the Concentration of Organic Acids

Figure 1 shows the powder XRD diffractograms after 20 h of synthesis of the dried gel with and without succinate added. When no succinate is added, an amorphous gel is observed as shown by the lack of XRD reflections and presence of an elevated background in the 2θ range 15-35° (Fig 1A), which does not possess any features from the smectite group of 2:1 layer silicates. In contrast, the gel with succinate added at concentrations of 0.01, 0.10, 0.50, and 1.00 M (Figure 1) shows an XRD peak emerge with a low-angle shoulder between 3.3 and 8.3° centered at 5.82° corresponding to a first-order basal reflection, *d*_0 0 1_ of sauconite. The *d*_0 0 1_ values, which corresponds to the thickness of the 2:1 phyllosilicate layer to layer distance (per half unit cell), increase for larger [succinate], varying between 14.2 and 15.1 Å (Figure 1B). In addition to the (0 0 1) reflection labeled in each XRD diffractogram, six more appear at 2θ angles of 11.46 (7.72 Å), 19.68 (4.51 Å), 28.60 (3.12 Å), 33.96 (2.64 Å), 59.98 (1.54 Å), and 70.20° (1.34 Å) most likely representing the (0 0 2), (0 2 *l*), (0 0 5), (1 1 0), (0 6 0), and (2 2 0) basal reflections of sauconite (ICDD PDF No. 00-008-0445),^16,17^ respectively. Interestingly, the weak peak at 4.51 Å that should correspond to the 0 2 *l* reflection suggests the sauconite is irregular and not perfectly oriented.^18,19^ The peak (0 6 0) at 1.54 Å confirms the trioctahedral structure of sauconite. The reversal in the trend of *d*_0 0 1_ with concentration above certain high [formate], or [acetate], [oxalate], or [malate] (Figure S1-S4) is associated to larger particle sizes limiting the process of expansion.^20^

The higher *d*^0 0 1^ basal spacings for the series of sauconite in Figure 1A with increasing succinate levels coincides with the growing total surface area (TSA) (Figure 1B). The larger TSA with increasing [succinate] is likely attributed to greater formation of the 2:1 layered structure. It is also probable that succinate has accumulated in the interlayer space, as supported by both the increase in *d*^0 0 1^ basal layer to layer distance and the lower than expected total surface area of a 2:1 layered sauconite. The highest TSA value in the experiment of Figure 1 is 338.8 (± 4.1) m^2^ g^-1^ (Figure 1B) indicates we are observing the beginning of the crystallization process. For comparison, the typical values of TSA for pure 2:1 layered clay minerals of the smectite family^6^ ranging from 600 to 800 m^2^ g^-1^ are only reached after 1 week of synthesis.

DRIFT spectra of the precursor gel, sauconite synthesized with [succinate] = 0.010, 0.10, 0.50, and 1.0 M, and sodium succinate are shown in Supplementary Fig. S5 online. In the gel, there is a broad band in the range 3100-3600 cm^-1^ corresponding to overlapping OH stretching bands due to water and the weaker corresponding OH-bending mode of water at 1645 cm^-1^ (Figure S5, dashed line).^19^ In addition, there is a broad stretching band near 1000 cm^-1^ in the precursor gel assigned to Si-O. After addition of succinate, two new shoulders appear in the spectrum at ~3639 and ~3743 cm^-1^, which are characteristic stretching vibrations of hydroxyl groups (*v*_OH_) attached to octahedral Zn ions located in the interior blocks of sauconite.^21^ IR bands appear at 2951 and 2973 cm^-1^ in the presence of succinate and these are assigned as C-H stretching vibrations of succinate, which match the spectrum of the sodium succinate standard.

The new band at 1630 cm^-1^ in sauconite (Supplementary Fig. S5 online) quickly grows from the stretching at 1645 cm^-1^ in the gel, due to OH bending vibrations of interlayer water.^18, 19^ This band at 1630 cm^-1^ due to the OH-bending mode of interlayer water is typical of 2:1 layered trioctahedral clay minerals.^22^ After addition of succinate, the bands from the amorphous gel at 604 and 1009 cm^-1^ show the prominent development of sauconite shoulders from symmetric Zn-O vibration at 661cm^-1^, and asymmetric in plane Si-O-Si vibration at 1020 cm^-1^.^20,23^ The bands at 1404 and 1383 cm^-1^ in the amorphous gel are due to a small amounts of sodium nitrate left from the synthesis.^24^ Spectra of sauconite also show bands at 1579 and 1296 cm^-1^, corresponding to the stretching of C=O and C-O groups in the structure,^17^ which are matched to sodium succinate standard (Supplementary Fig. S5 online). Interestingly, the absorbance from succinate incorporated in the interlayer is correlated to the amount added during the synthesis. The intercalation of aliphatic organic carbon in 2:1 layered clay minerals has been reported in pure standards and natural clays extracted from sediments.^25,26^ Overall, two analytical methods (XRD and DRIFT spectroscopy) confirm a clay mineral is successfully synthesized in the presence of central metabolites such as acetate, malate, and succinate.

## Effects of Temperature and pH

Figure 2 shows the powder XRD diffractograms for 20 h syntheses with 0.10 M succinate under variable reflux temperature. While at 70 °C no sauconite formation is registered, the spectroscopic features of the mineral are observed at 75 °C. However, the first-order reflection (0 0 1) is only well-resolved for a minimum temperature of 80 °C, which allows the measurement of *d*_0 0 1_ displayed in Figure 2B. As the first-order reflection (0 0 1) becomes better defined with increasing temperature, its width decreases indicating the layer to layer distance expands. For example, *d*_0 0 1_ increases in Figure 2 from 13.6 Å at 80 °C to 14.2 Å at 90 °C. Control experiments demonstrate that sauconite dried overnight under room temperature (Supplementary Fig. S6 online) showed the same XRD features observed when drying at 90 °C (Figure 2) was applied. Related experiments changing the pH of synthesis for 0.10 M succinate, under reflux at 90 °C for 20 h, demonstrate (see *d*_0 0 1_ values in Supplementary Fig. S7 online) that the best crystallinity is obtained at pH_0_ 9.0. Summarizing, the best conditions for studying the time series of the synthesis by microscopy techniques correspond to 1.0 M succinate, pH_0_ 9.0, and 90 °C for reflux. Comparison of samples after 20 h of synthesis under these optimized conditions, which have been Mg-saturated first and glycerol solvated second, revealed a 3.4 Å shifting of the (0 0 1) reflection (Supplementary Fig. S8 online). Thus, expansion behavior for a double layer trioctahedral smectite is confirmed to be in the range of reported values (3.1-3.7 Å) for 2:1 layer smectites.^27^ The unexpected lower intensity observed after glycerol saturation confirms the 2:1 layer sauconite lacks any stacking of layers at 20 h.

**Figure 2.**
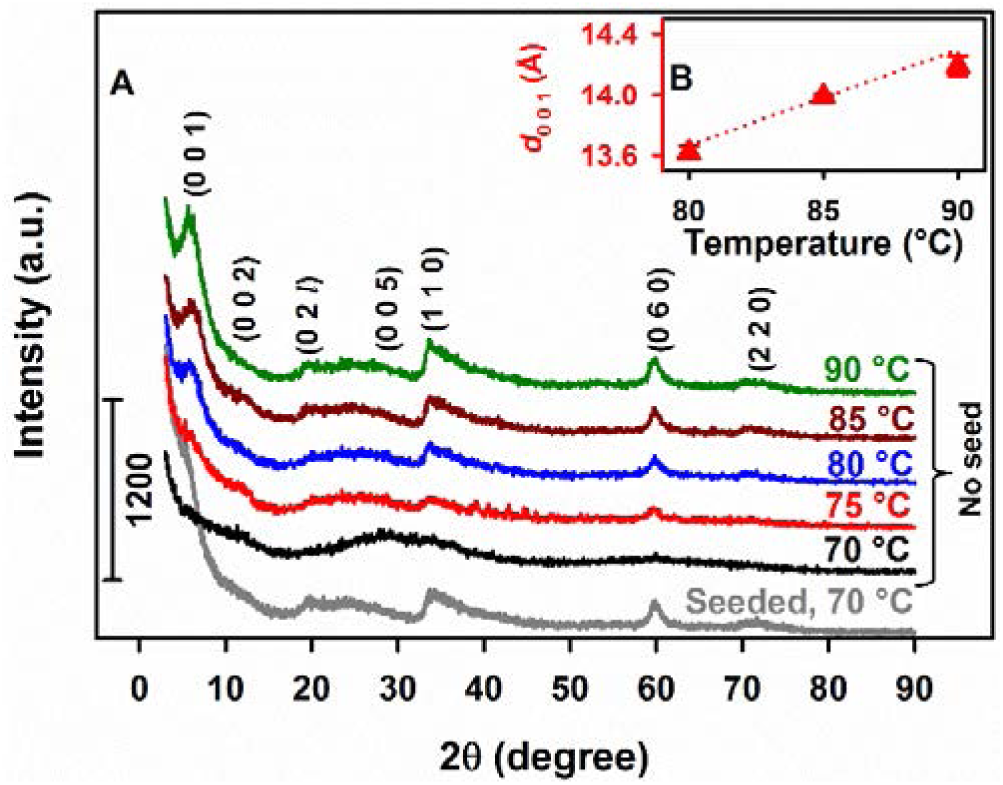
Powder XRD and *d*_0 0 1_ values for 2:1 sauconite synthesized at pH_0_ 6.3 with 0.10 M succinate during 20 h under variable temperature. (A) XRD diffractograms under the temperatures listed above each trace. The gray line at the bottom labeled “seeded” corresponds to an experiment at 70 °C started after adding a single particle from sauconite synthesized at 90 °C. An oval shaped particle weighing 0.7 (± 0.1) mg and symmetry axes of length 313 (± 9) and 144 (± 7) μm was used for seeding. (B) Values of *d*_0 0 1_ (red solid triangle) for traces in (A) with no seed vs temperature.

## Seeding Induced Crystallization

The catalytic power of the synthesized clay is confirmed by repeating the procedure at 70 °C after spiking the gel with a single (macroscopic) sauconite particle obtained at 90 °C. As shown by the bottom trace labeled “seeded” in Figure 2, after spiking the single particle, the synthesis is promoted at 70 °C in the same 20 h timespan, yielding 1.86 (± 0.06) g of sauconite (a 54% of the material obtained at 90 °C). The fact that a simple particle of sauconite serves as a seed crystal to growth a larger amount of mineral serves as an outstanding example of the self-catalytic power of clays toward their crystallization. The surface of the seed particle facilitates heterogeneous nucleation at lower temperature by reducing the activation energy for crystallization. The observed acceleration in crystallization by seeding does not rely solely on random events but results from the surface interaction with chemical species in the precursor gel at 70 °C. The seed particle interacts with soluble free and complexed ions moving randomly in the gel, establishing intermolecular forces that are needed to form the crystal lattice. The addition of a seed particle to the gel provides a route to direct a process that does not depend on random interactions anymore.

## Morphological and Structural Analysis

Complimentary preparation, imaging and analytical techniques were applied to investigate the evolution of sauconite synthesized with 1.0 M succinate at pH_0_ 9.0 and 90 °C. Samples at the early (0 h), intermediate (6 h) and final stage (20 h) of the synthesis were selected for detailed analysis. The sample at zero timepoint revealed the presence of unstable Zn-containing plate-like particles, likely Zn(OH)_2_, that are coated with Zn-containing nanoparticle, possibly of zinc silicate (Zn_2_SiO_4_) (Supplementary Fig. S9 online). However, the previous phases were only observed in the untreated, air-dried condition for TEM and were likely unstable under the conditions required for plastic embedding or cryo-fixation. TEM images and energy dispersive X-ray spectroscopy (EDS) demonstrate that the morphology of the precursor gel corresponds to an amorphous material that has not incorporated aluminum in the structure (Supplementary Fig. S9 online).

After 6 h of synthesis the sample showed aggregates of sauconite nanocrystals, sometimes in association with an unknown gel-like phase (Figure 3A–C), and occasionally Zn-containing nanocrystals as seen in the sample at the zero timepoint. These latter crystals have an atomic structure close to Zn^2^SiO^4^.with lattice fringes spaced by ~2.3 Å (Figure 3C). Sauconite is made up of Al, Si, H and O in ratios similar to other smectite-group minerals (Figure 3D). Compared to the Zn-clay synthesized, the Zn-containing particles contain more Zn, much less Si and no Al. These nanoparticles were not observed in the TEM images from samples prepared for ultrathin section suggesting they are sensitive to the chemical treatment required for sample processing i.e. dilution, washing, dehydration, embedding. The objective of the TEM analysis of the samples in ultrathin section was to image the lattice fringes and observe the stacking order of individual 2:1 layers. High-resolution TEM (HRTEM) analysis of the cross-section through aggregates of sauconite showed disorganized and random arrangements of individual 2:1 layers. Most smectite-group minerals, including synthetic saponite ^13^, consist of packets of short-range, coherent stacks of 2:1 silicate layers. Both the structure and chemical composition of sauconite resembles, however, other known smectite-group minerals.

**Figure 3.**
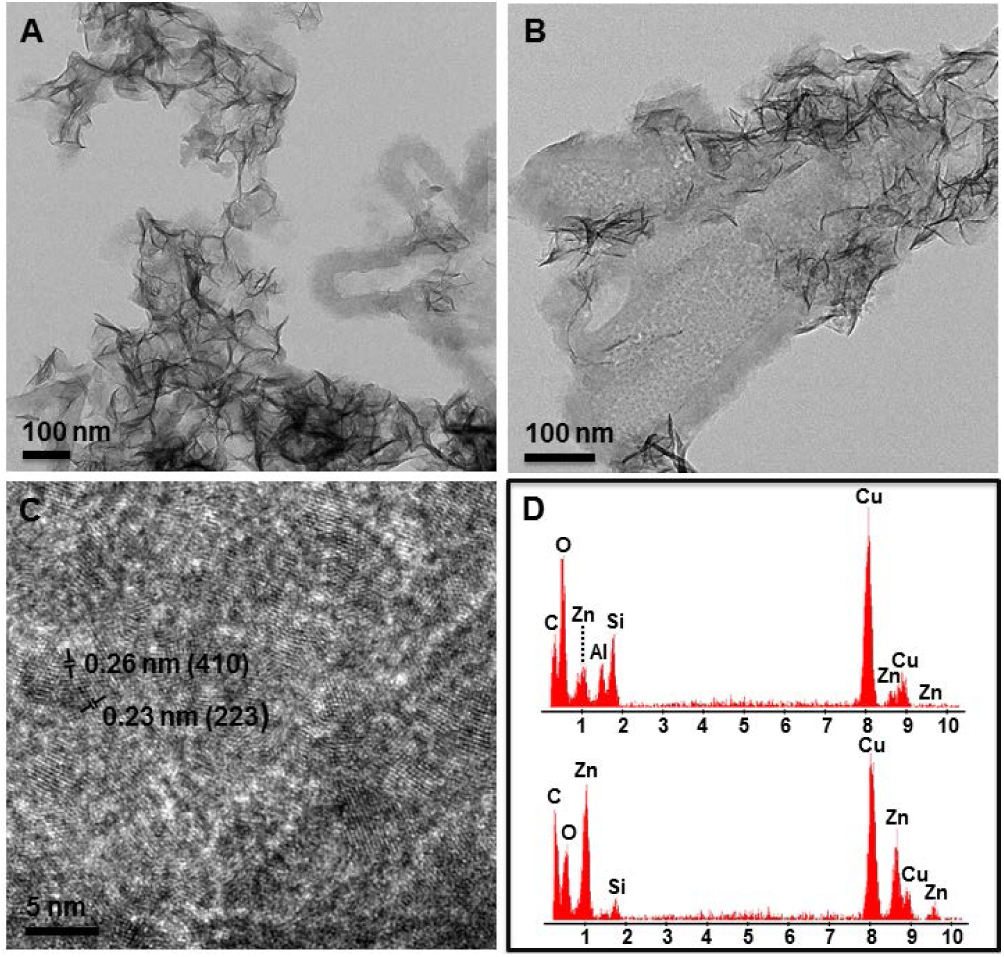
TEM images and energy dispersive X-ray spectra (EDS) of air-dried whole mount synthesized sauconite with 1.0 M succinate at pH_0_ 9.0 after 6 h reflux at 90 °C. (A) Aggregates of sauconite nanocrystals. Gel-like material shows nucleation of sauconite (right side of image). (B) Higher magnification of the gel-like sauconite showing the presence of unknown zinc-containing nanocrystals in addition to sauconite. (C) Lattice fringe image of zinc nanoparticles. (D) Representative EDS spectra of the composition of sauconite nanocrystals (upper) and zinc nanoparticles (lower).

The structure and chemical composition of the sauconite 2:1 layers after 20 h of synthesis is the same as the sample collected after 6 h. There is, however, a difference in the organization and aggregation of individual 2:1 layers as well as in the distribution and density of the particles. The TEM images for the 20 h synthesis in ultrathin section shows only isolated stacks containing a few individual 2:1 layers of sauconite (Figure 4). The corresponding XRD diffractogram for this sample shows the 2:1 layer to layer distance expands a rate of 1.54 Å day^-1^, which is accompanied by the incorporation of Al. Aluminum incorporation could take place by a 3 Zn^2+^⇆ 2 Al^3+^ + 1 vacancy substitution mechanism, which would not introduce a positive charge to the octahedral sheet.

**Figure 4.**
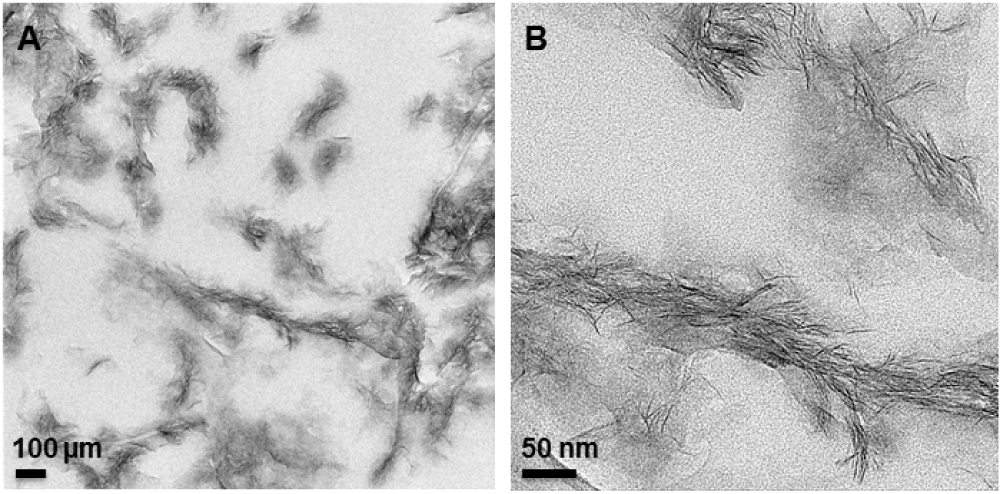
TEM images of ultrathin sections of synthesized sauconite with 1.0 M succinate at pH_0_ 9.0 after 20 h at 90 °C. (A) Overview of the area in the section containing aggregates of sauconite crystals. (B) Close-up of an aggregate of sauconite in the center of (A) showing the disorganized arrangement of individual 2:1 layer silicates with the occasional short-range stack of coherent layers.

### Electron Tomography and 3D Reconstruction

Electron tomography (ET) was carried out to determine the detailed 3D structure and mode of interaction of the sauconite 2:1 layers in ultrathin sections. The objective in ET is to reconstruct the 3D structure of an object from a series of 2D projections. Using the single-axis tilting method, a series of TEM images was collected from isolated aggregates of sauconite 2:1 layers in ultrathin section at every 2º tilt angle from −70º to +70º. No significant differences were observed between samples after 6 and 20 h of synthesis. A snapshot of the tomogram and its reconstructed image are shown in Figure 5A–B. The images show the individual 2:1 layers are arranged in a random edge-to-face orientation within isolated aggregates.

**Figure 5.**
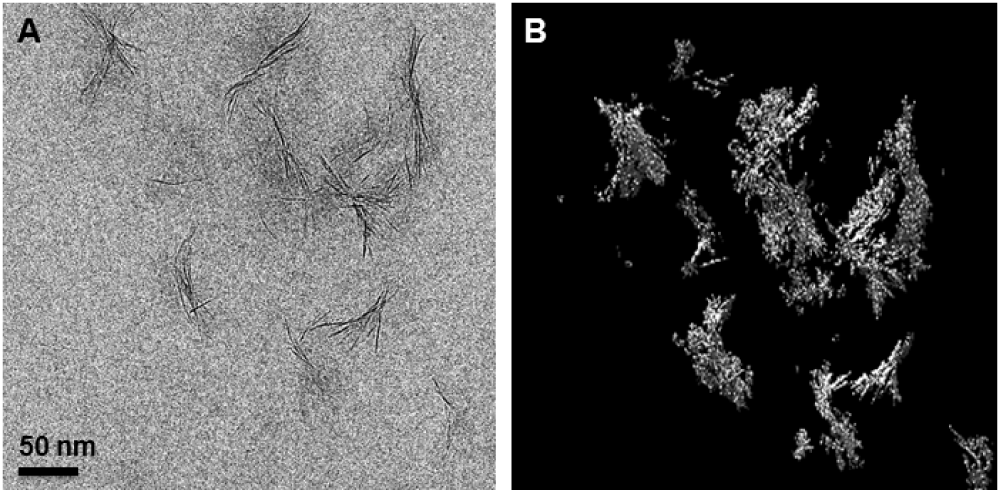
(A) Snapshot from a tomogram of an isolated aggregate of sauconite nanocrystals from a less dense area in the section for 20 h synthesis with 1.0 M succinate at pH_0_ 9.0 and 90 °C (B) 3D reconstruction of (A). See *Movie* M4 in the *Supporting Information*.

Cryo-TEM was applied to study the size and shape of individual sauconite particles and their distribution in their natural dispersed state. The results of the extremely diluted (0.2 wt %) frozen samples show significant differences between samples collected after 6 and 20 h of synthesis (Figure 6). Figure 6A shows the emergence of the sauconite particles from the gel-like precursor suggesting initial nucleation and growth of the sauconite phase. Surprisingly, the sauconite 2:1 layers were not dispersed in either sample confirming the features and morphology observed in the ultrathin sections (Figure 5) is not an artifact, but a growth feature. Figure 7 displays snapshots from the tomograms of the sample after 20 h of synthesis (see Supplementary *Video* M1 online). Figure 7C–D shows the aggregation of individual sauconite particles consists of stacks of a few semi-oriented 2:1 layers suggesting a unique growth feature. Most synthetic or natural smectite-group minerals are completely dispersed as individual particles consisting of single 2:1 layers or stacks of a few ordered face-to-face 2:1 layers.^28^ Different rotational views are displayed in Supplementary *Videos* M2 and M3 online.

**Figure 6.**
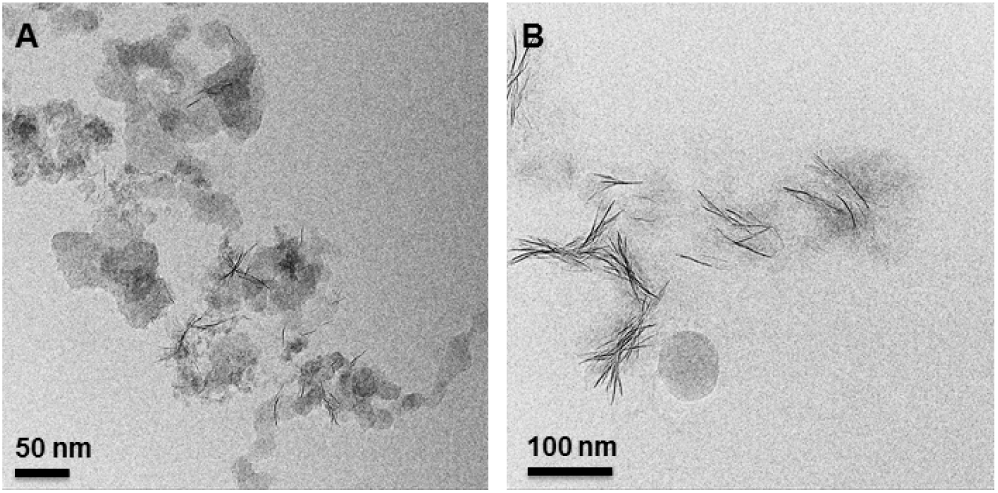
(A) Cryo-TEM image of sauconite synthesized with 1.0 M succinate at pH_0_ 9.0 after 6 h at 90 °C. The presence of irregular-shaped, possibly gel-like particles seen in Figure 3A-B in their dispersed state. The initiation of the formation of sauconite nanocrystals within these particles can be seen in the center of the image. (B) Cryo-TEM image of 20 h synthesis as for Figure showing isolated, disorganized aggregates of sauconite nanocrystals also seen in Figure 4. This sample in a diluted dispersed suspension consisting of pure sauconite nanocrystals.

**Figure 7.**
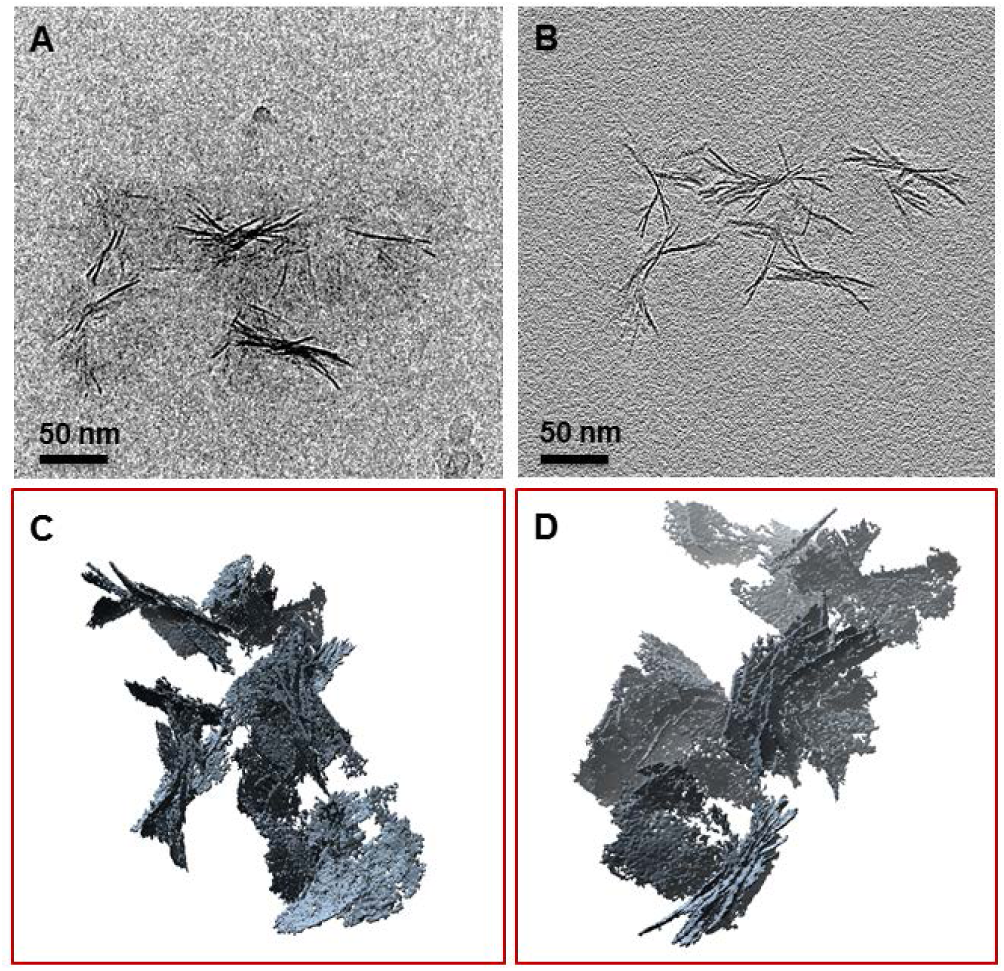
Snapshots from tomograms and 3D reconstructions for 20 h synthesis with 1.0 M succinate at pH_0_ 9.0 and 90 °C from cryo-TEM. (A) Raw image showing the individual aggregates of sauconite nanocrystals. (B) Reconstructed image of (A). (C) (D) Different views of reconstructed aggregates of sauconite nanocrystals seen in (A) and (B) showing the 3D arrangement and orientation of individual nanocrystals. See *Movies* M2 and M3 in the Supporting Information.

### Kinetics of Crystallization and Swelling

The results presented above confirm the formation of sauconite by several complementary methods. While TEM and cryo-TEM provide images of sauconite, they both indicate that XRD and DRIFT spectroscopy can be used to monitor the time series of Zn clay formation. Thus, a series of syntheses under the optimized conditions of 1.0 M succinate, pH_0_ 9.0 at 90 °C were performed and stopped at 0, 1, 2, 6, 15, and 20 h to monitor sauconite formation. Figure 8A and B show the timepoints for XRD diffractograms and DRIFT spectra. The (0 0 1) peak becomes evident after only 1 h and undergoes a pronounced broadening at 20 h. Similarly, the stretching vibrations for sauconite are clearly registered in the time series of Figure 8B, what makes possible to estimate a time profile for the incorporation of succinate. Finally, the time series of measured TSA values for the corresponding samples is displayed in Figure 8C, where *d*_0 0 1_ values extracted from Figure 8A and the integrated area under the C-H stretching (ν_C-H_) between 2937 and 2991 cm^-1^ from Figure 8B are also presented. Remarkably, the expansion of the layer to layer distance (*d*_0 0 1_), the total surface area, and the amount of succinate accumulated in the interlayer, all depend on time with a similar exponential correlation (Figure 8C).

**Figure 8.**
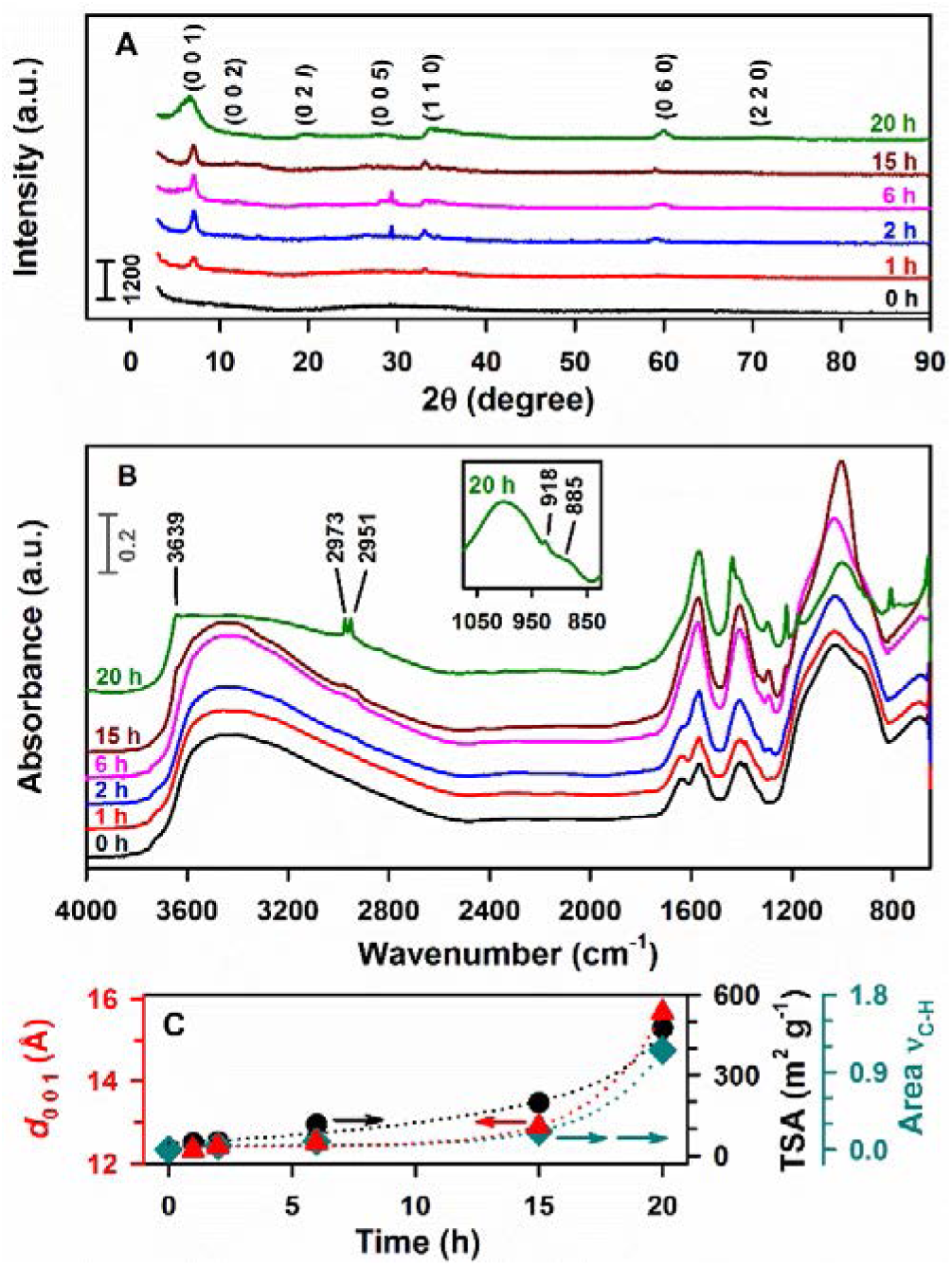
Time series of powder XRD diffractograms and DRIFT spectra registered at 0, 1, 2, 6, 15, and 20 h for sauconite synthesis with 1.0 M succinate, pH_0_ 9.0 at 90 °C. (A) XRD diffractogram. (B) DRIFT spectra including an inset with features at 20 h. (C) Time correlation of *d*_0 0 1_ values for 2:1 sauconite (red solid triangle) in (A), TSA (black solid circle) in (A), and the area for C-H stretching (*Ȟ*_C-H_, teal solid diamond) integrated between 2937 and 2991 cm^-1^ in (B), after baseline correction with a two point algorithm.

The peak at 918 cm^-1^ in the DRIFT spectra of synthesized sauconite at pH_0_ 9.0 from 0 to 6 h (Figure 8B) corresponds to Al-OH bending from the reactant 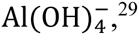 the dominant hydroxy complex of aluminum in the starting gel at high pH.^14^ As the previous peak disappears after 15 h of synthesis, a new peak is developed at 885 cm^-1^ assigned to Si-O-Al^IV^ vibrations in the tetrahedral position of sauconite. The stronger intensity for the Si-O-Al^IV^ vibration at 20 h in Figure 8B, which is also present in experiments with variable [succinate] (Supplementary Fig. S5 online), suggests the incorporation of Al to the tetrahedral layer,^22^ which is key for crystallization. The role of succinate as a catalyst is proposed to facilitate the simultaneous insertion of two Al^3+^ above and below a plane that becomes the interlayer of sauconite. Figure 9 displays a representation that takes into account the previous concept for the layers in 2:1 trioctahedral sauconite, which agrees with cryo-TEM, XRD, and DRIFT spectroscopy observations. In this model representation, succinate and the water molecules are slightly scaled-up relative to the centers in the crystalline structure to facilitate their visualization. The constrains that succinate experience to accommodate in the interlayer space is thought to be controlled by the slower free rotation of σ-bonds among carbon atoms in the gel compared to the freedom provided in a complete solution state. The 3D structure of succinate is represented in Figure 9 for the staggered conformation that decreases the torsional strain in energy from eclipsing interactions by optimizing the rotation around carbon-carbon σ-bonds. Because *d*_0 0 1_ grows over time (Figure 8C), it is logic to assume that the sauconite structure in Figure 9 reaches its maximum swelling as the conformer of succinate optimizes its geometry reaching a length of 5.6 (± 0.7) Å. In addition, the selected succinate structure appears stabilized by interactions with two aluminum centers located on opposite sites of the interlayer space (Figure 9). However, it would also be possible to envision interactions of succinate with a single zinc center of the interlayer. Figure 9 also depicts the tetrahedral–octahedral–tetrahedral sheets with Zn, Al and Si, as proved by EDS (Figure 3D).

**Figure 9.**
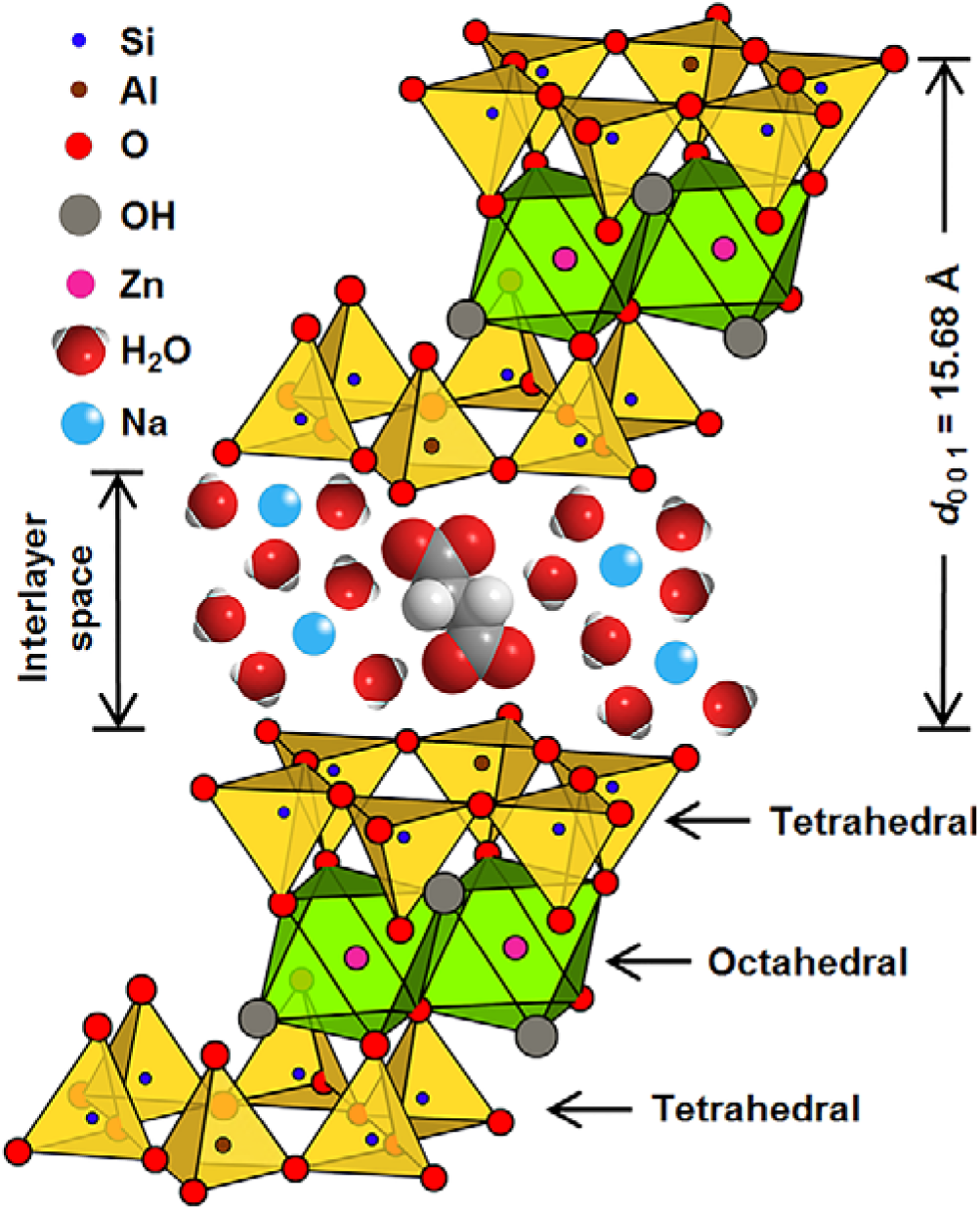
Model displaying the stacking order of layers in a 2:1 trioctahedral sauconite structure al synthesized during 20 h using 1.0 M succinate at pH_0_ 6.3 and 90 °C. The layer to layer distance *d*_0 0 1_ = 15.68 Å is represented to be about 3-times larger than the length of succinate that appears longitudinally oriented perpendicular to tetrahedral aluminum centers.

The peaks at 1438 and 1570 cm^-1^ (Figure 8B) at 20 h of synthesis represent symmetric and asymmetric stretching vibrations for C=O (*ν*_C=O_) in the carboxylate group of succinate, respectively.^17^. In addition, the two undissociated -COOH groups of succinic acid (pK_a1_ = 4.21 and pK_a2_ = 5.64)^30^ expected at 1297 cm^-1^ (*ν*_C=O_) and 1691 cm^-1^ (*v*_C-OH_)^31^ are absence at pH_0_ 9.0 because only completely dissociated succinate is available in equilibrium. However, at pH_0_ 6.3 the fraction of dianion drops to 81.9% while succinate monoanion reaches an 17.9% justifying the observation of a peak at 1297 cm^-1^ for the symmetric *v*_C-OH_ stretching of –COOH groups in Figure 8B. Therefore, DRIFT spectroscopy confirms the dianion of succinate depicted in Figure 9 is the dominant species both participating in the synthesis and accumulating in the interlayer space of sauconite in experiments at pH_0_ 9.0. Total surface area measurements, TEM and cryo-TEM together with XRD and DRIFT spectroscopies reveal that the slow initial swelling during sauconite growth becomes exponentially faster after aluminum incorporation to the structure. The total surface area of the synthesized clay is a direct function of the amount of a simple dicarboxylic acid, which is an adsorbate in the interlayer space (Figure 9) and a catalyst for crystallization. Therefore, as the amount of organic salt increases, swelling occurs and the reactive surface growths exponentially. The associated retention of water in the interlayer space is enhanced for longer syntheses as indicated by the 2.2 × 10^3^ expansion in TSA occurring when transitioning from 0 to 20 h for the data in Figure 8C.

## Conclusions

Because clay minerals contribute most of the inorganic surface to catalyze reactions in soils, these results provide new understanding of the continuous evolution of present agricultural fields. Overall, this laboratory study demonstrates an effective coupling between photochemistry mediated by semiconductor minerals and clay formation exists. The work also demonstrates the self-catalytic power of clays toward mineral formation in the geological past as well as the ability of simple central metabolic molecules to co-catalyze crystallization in short time scales. The co-catalytic role of succinate between layers is to provide soluble complexes of Al^3+^, which is the limiting reagent for sauconite crystallization. Once the likely intercalation of succinate (or the other organic species) balances the relative adsorption forces between the precursor gel and the aluminum species, due to its bidentate ability, it can overturn the attraction forces between adjacent sauconite whiskers. A detailed description of the nucleation and growth of model clay particles is supported by cryogenic and conventional TEM. Finally, a major outcome of this work is that photochemistry may have played an important role in the origin of life on early Earth and other rocky planets.

## Methods

All syntheses are performed by duplicate. The analyses by the methods listed below are reported as mean values and standard deviations (± SD) from independent experiments.

### Preparation of Sauconite

The synthesis of Na^1.2^Zn^6^ [Si^6.8^Al^1.2^]O^20^(OH)^4^•*n*H^2^O started by diluting 4.0 g of Na^2^SiO^3^ solution (26.5 Wt. % SiO_2_) in 10 mL water. A solution of 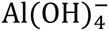 was prepared by dissolving 1.21 g of Al(NO_3_)_3_•9H_2_O in 8 mL of 2.0 M NaOH. The addition of 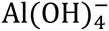 solution to pre-diluted Na_2_SiO_3_ under continuous stirring, produced a cloudy gel that was left standing without stirring for 1 h before further use. The gel was stabilized with 1 mL of concentrated HNO_3_ before addition of 4.81 g of Zn(NO_3_)_2_•6H_2_O dissolved in 100 mL water. The synthesis was performed under reflux at 90 °C,^15^ unless noted otherwise, by augmenting the gel with the sodium salt of succinic acid to reach a concentration in the range 0.010–1.0 M. Alternatively, formic acid, acetic acid, oxalic acid, or L-malic acid sodium salts were also employed (Supplementary Information). After 1 h of adding the organic salt, 2.0 M NaOH was used to vary the pH of gels with 0.10 M succinate in the range 6.4–13.9. The weight of sauconite obtained after 20 h under these conditions (0.10 M succinate at pH^0^ 7.0 and 90 °C) was 3.43 (± 0.01) g. In addition, for 0.10 M succinate, 20 h syntheses under stirring with reflux temperature from 70 to 90 °C every 5 °C were performed. Finally, the sauconite was centrifuged (5 min at 4400 rpm), triple washed with water, and dried overnight at 90 °C, unless noted otherwise.

### Powder XRD Diffractograms

A D8 Advance Bruker AXS diffractometer (Cu Kα *λ* = 1.5418 Å) provided crystal structures. The diffractogram of samples mounted onto the sample holder was recorded from 2° to 90° at a scan rate of 0.01° s^-1^. The distance of the 2:1 layer (*d*_0 0 1_) was calculated using Bragg’s law: *d*_0 0 1_ = *nλ*/(2 Sin*θ*), where *λ* = 1.5418 Å (Cu Kα radiation) is the wavelength of X-ray, *θ* is the scattering angle, and the integer *n* is the order of the corresponding reflection. A traditional DIFFRAC method was applied to subtract the sharply increasing scattering of the incident beam from the background at low angle.^23^ The method corrects the background by optimizing its maximum concavity, and enables the calculation of accurate *d*_0 0 1_ values, e.g., 15.10 ± 0.09 Å. The application of this method is checked by measuring *d*_0 0 1_ = 15.46 Å for a saponite standard (Mg_2_Al)(Si_3_Al)O_10_(OH)_2_•4H_2_O from the Cuero Meteorite Crater in DeWitt Co. Texas (Excalibur Mineral Company). The diffractogram of the standard matches the pattern of saponite-15Å (ICDD PDF No. 00-030-0789).^17^

The preparation of oriented sauconite slides for XRD analysis in Figure S7 (Supplementary Information) was based on Drever’s method.^32^ Briefly, 25 mL of 0.50 M MgCl_2_ was mixed in a 50 mL centrifuge tube with 200 mg of sauconite synthesized during 20 h with 1.0 M succinate at pH_0_ 7.0 and 90 °C. After 1 min sonication and 5 min centrifugation (3000 rpm), the supernatant was discarded. This washing procedure was repeated by triplicate before a final wash with DI water to remove extra Mg^2+^. The unflocculated suspension was filtered (0.45 μm pore size and 47 mm diameter) under vacuum. The sauconite sample was carefully transferred onto a glass slide (26 × 46 mm) as described in the filter-membrane peel technique.^32^ The Mg-saturated glass slides were analyzed by XRD from 2 to 40° 2*ș* degrees with a step scan of 0.07° (2*ș*) and a time per step of 4 s. Finally, the Mg-saturated sample was exposed to a vapor saturated glycerol atmosphere in a desiccator for 24 h before scanning by XRD. While the 2:1 layer structure is confirmed, the reversal in the intensities expected for both samples suggests there is no oriented stacking of layers, explaining the difficulties encountered during sample preparation.

### DRIFT Spectra

DRIFT spectra were recorded (200 scans) with a Nicolet 6700 FTIR spectrometer and analyzed with OMNIC32 software (both Thermo Fisher Scientific). A liquid N2 cooled MCT detector was employed in the range 600-4000 cm^-1^ with 4 cm^-1^ resolution. Oven dried samples of synthesized sauconite (30 mg) were homogenized with spectroscopic grade KBr (500 mg), and then poured into the smart collector diffuse reflectance accessory, which optics were continuously purged with N_2_(g). The incorporation of succinate over time was monitored by integrating the area under the C-H (*Ȟ*_C-H_) between 2937 and 2991 cm^-1^.

### Measurement of TSA

TSA is determined using the ethylene glycol monoethyl ether (EGME) method^33^ for 0.5 g of sauconite synthesized 1) under variable [succinate] at pH_0_ 6.3, and 2) for pH_0_ 9.0 and 0.10 M succinate during the 20 h time series. Samples and a kaolinite standard (0.5 g each) were oven dried (90 °C) overnight, and analyzed following the standard procedure reported in the literature.^33^

### TEM, EDS, and Cryo-TEM

To image particle size and shape of the samples, aqueous suspensions (5 μL) from the synthesis were dispersed and pipetted onto a 200-mesh copper TEM grid with carbon support film (whole mount), and allowed to dry under air at room temperature. For high-resolution TEM (HRTEM), sampled from synthesis at different timepoints were embedded in EPON liquid epoxy resin. Approximately 10-20 mg of the sample was placed into 1.5 mL polypropylene Eppendorf micro-test tubes and dehydrated by adding 100% ethanol to remove adsorbed water. After centrifugation at 13000 rpm for 15 min, the supernatant was removed and the sample dispersed in a mixture of 10% EPON resin and 90% ethanol by ultrasonic treatment. These steps were repeated with mixtures containing 30, 50, 70, and 100% EPON.

To obtain complete dispersion of the material, the ethanol:resin mixtures were agitated in an ultrasonic bath for ~1 min. Each incubation step lasted 24 h with the samples placed onto a rotator to ensure continuous agitation. The resin-clay mineral mixture was transferred into embedding moulds after the fifth incubation step (100% EPON resin). After a settling time of 1-2 h, the samples were polymerized by placing the moulds into an oven at 65 ºC for 48 h. Ultrathin sections (70-80 nm) were cut from the polymerized resin blocks using an ultramicrotome and transferred onto 200-mesh copper TEM grids with carbon support film. The samples on whole mounts and in ultrathin section were imaged with an FEI Tecnai G^2^ F20 200 kV TEM equipped with a Gatan Ultrascan 4000 CCD Camera System Model 895 and EDAX Octane T Ultra W /Apollo XLT2 SDD and TEAM EDS Analysis System.

For cryo-TEM, 5μL of the aqueous suspension of the synthesized sauconite (previous to the final drying and washing steps) was transferred onto a C-flat holey carbon sample support grid (R2/2; Protochips, Inc.). Excess fluid was blotted and the sample flash frozen hydrated by plunging into a bath of liquid nitrogen-cooled liquid ethane using the FEI Vitrobot Mk IV Grid Plunging System (FEI Co.). The grids were stored in liquid nitrogen until imaged in a FEI Titan Krios 300 kV Cryo-S/TEM equipped with a Falcon 2 direct electron detector (DED) (FEI, Inc). Images were collected at a magnification of 75k× corresponding to a pixel size of 1.41 Å and a defocus level ranging from −2.0 to −3.0 μm, under low dose conditions. Tomograms from the cryogenic and epoxy embedded samples were collected using FEI Batch Tomography Software version 4.0. The cryogenic tilt series was collected at a magnification of 59k× every 2º over a tilt range of ± 70º. The nominal pixel size was 0.14 nm with defocus of -2μm

### Image Processing

Images from the single-axis tomograms were aligned, filtered and reconstructed into a series of tomographic slices using IMOD (version 4.8.26).^34^ The back projection method was used for reconstruction. The 3D visualization and surface models were created with UCSF Chimera (version 1.10.1).^35^

## Associated Content

### Supplementary Information

Supplementary Figures S1-S9 and Videos M1-M4 available online.

### Author Information

**Notes:** The authors declare no competing financial interest.

## Acknowledgements

M.I.G. acknowledges funding from the National Science Foundation. Partial support from the University of Kentucky by a Research Challenge Trust Fund Fellowship to R.Z. is gratefully acknowledged. H.V. acknowledges funding from the Natural Sciences and Engineering Research Council (NSERC) of Canada.

